# FetchM: Streamlining Genome and Metadata Integration for Microbial Comparative Genomics

**DOI:** 10.1101/2025.04.08.647722

**Authors:** Tasnimul Arabi Anik

## Abstract

FetchM is a Python-based tool for fetching, analyzing, and combining bacterial genomic metadata from the NCBI Genome database and associated sample metadata from NCBI BioSample records. When working with bulk-genome analyses, such as comparative genomics or pangenome studies, you often require a unified dataset that captures the full context of a particular bacterial species population. You can obtain genomic metadata by downloading the ncbi_dataset.tsv file from the NCBI Genome database for a specific bacterial species. However, this file lacks key metadata fields such as Collection Date, Host, Geographic Location, and Isolation Source. FetchM fills this gap by automatically retrieving these missing fields, linking genome accessions to their corresponding BioSample records via the NCBI Entrez API. FetchM not only helps you compile a complete, metadata-rich dataset but also provides visualizations and summaries of both genomic and contextual metadata features. You can filter and download sequences based on specific criteria such as year, host, isolation source, country, continent, and subcontinent, making it a flexible and powerful companion for large-scale genomic studies. FetchM is available as an open-source tool at: https://github.com/Tasnimul-Arabi-Anik/FetchM. It can also be downloaded as a PyPI package.

## Introduction

Accurate and complete metadata associated with genomic datasets are critical for downstream bioinformatics analyses, comparative studies, and data reuse in microbiome research [1]. Public repositories such as the National Center for Biotechnology Information (NCBI) provide a wealth of microbial genomic data through platforms like BioSample and GenBank [2]. While GenBank contains data related to genome sequences, BioSample provides metadata describing the biological source of those genomes [3]. Both types of information carry significant value for understanding the genomic context and drawing meaningful biological interpretations, especially in large-scale studies such as pangenome or comparative genomic analyses [4]. However, retrieving and integrating both datasets can be time-consuming and technically challenging, especially when dealing with large genome collections [5].

NCBI provides tools that assist researchers in retrieving critical genomic and metadata [6]. Among them, the NCBI E-utilities (Entrez Utilities) is a powerful suite of programmatic tools that allows users to access and retrieve data from various NCBI databases, including GenBank, BioSample, and PubMed [7]. These utilities support automated querying, downloading, and parsing of large-scale biological data via HTTP requests, streamlining data acquisition for bioinformatics workflows [8]. However, retrieving the biological source metadata for genomes using these tools can still be time-consuming. Having a combined dataset that integrates both genomic data and associated metadata would greatly facilitate downstream analyses and enhance biological interpretation.

FetchM (https://github.com/Tasnimul-Arabi-Anik/FetchM) addresses these challenges by automating the retrieval, quality control, and visualization of genomic metadata from NCBI BioSample records. FetchM supplements incomplete metadata in standard NCBI downloads (e.g., ncbi_dataset.tsv) by programmatically querying BioSample APIs, filling gaps in fields such as collection date, geographic origin, host species, and isolation source. This contrasts with tools like EDirect [7], which require manual query construction. By enforcing user-defined CheckM completeness thresholds (default: 95%) and filtering based on taxonomy status, FetchM ensures dataset integrity—a feature absent in generic metadata fetchers like pysradb [9]. FetchM generates publication-ready visualizations (e.g., temporal trends, geographic distributions) without requiring coding expertise, bridging a gap left by script-heavy tools like ggplot2 [10].

This tool streamlines workflows for researchers requiring high-quality, annotated datasets for downstream analyses like phylogeography, host adaptation studies, or temporal trend assessments. Unlike NCBI’s standalone datasets tool, FetchM enables simultaneous metadata curation and genome sequence downloading, filtered by criteria such as host species or collection year. Besides, FetchM can annotate geographic location based on continent and subcontinent regions and allow users to download sequences from a specific region [11].

## Methodology

### Fetching Input Files

FetchM begins by processing an input file—typically ncbi_dataset.tsv—obtained from NCBI Datasets or genome assembly downloads. This file often lacks key contextual metadata such as collection date, geographic location, host, and isolation source.

### Metadata Retrieval from NCBI

FetchM uses the BioSample Entrez API via Biopython’s Bio.Entrez module to fetch metadata for each sample (12). It parses the XML structure of BioSample records to extract key fields and appends them to the original dataset.

### Quality Filtering

FetchM incorporates a multi-layered quality control process to select high-quality genomes for downstream analysis. The pipeline uses CheckM to assess genome quality metrics including completeness and contamination, with configurable thresholds (commonly ≥90% completeness and ≤5% contamination) (13). Filtering also incorporates ANI-based taxonomy check results.

### Data Processing and Standardization

The metadata curation step leverages the pandas library for data cleaning and standardization (14). This includes removal of duplicates, handling of missing fields through imputation or flagging, and consistent formatting of metadata categories such as taxonomic lineage and geographic location.

### Metadata Analysis and Visualization

FetchM supports metadata-driven analysis by generating summary statistics and frequency distributions for variables like host taxonomy, isolation source, location, and collection time. For visualization, it utilizes matplotlib (15), seaborn (16), and plotly (17) to create taxonomic bar plots, quality histograms, and interactive geospatial maps.

### Data Export and Integration

FetchM facilitates downstream analysis by exporting curated datasets in CSV/TSV formats compatible with tools like Roary (18), Panaroo (19), and anvi’o (20). Users may also retrieve complete genome sequences directly via NCBI Datasets CLI integration, ensuring compatibility with pangenomic workflows.

### Validation

To validate FetchM, we retrieved all available genomes and associated metadata for *K. oxytoca*. The genomes were filtered based on quality metrics including assembly completeness and ANI taxonomy check before proceeding with downstream analyses.

## Results and Discussion

To validate the functionality of FetchM, we applied the tool to the genomic dataset of *Klebsiella oxytoca*. As of April 7, 2025, a total of 1,304 genomes of *K. oxytoca* were available in the NCBI

Genome database. The genomic metadata was downloaded in tabular format and enriched using FetchM by retrieving additional sample-level metadata from the NCBI BioSample database.

FetchM was installed in a separate Conda environment and run using default parameters, which included a CheckM completeness threshold >95%, taxonomy status set to “OK,” and a 0.5-second delay between requests to comply with NCBI API guidelines (21). Out of the 1,304 genomes, 707 lacked CheckM values and were excluded from quality filtering by default. The remaining 597 genomes satisfied the taxonomy criterion and had completeness scores; among these, 524 genomes passed the strict CheckM filtering. Users can optionally disable this filter or run local CheckM analysis if they wish to retain genomes without embedded completeness data.

Among the 524 high-quality genomes, total sequence length ranged from 5,404,074 to 6,859,794 bp, with a mean of 6.09 Mb and a median of 6.06 Mb. The protein-coding gene counts varied from 4,951 to 6,635, with an average of 5,653 genes per genome, while pseudogene counts ranged from 31 to 300, averaging approximately 105. The distribution of these variables are shown in Figure 1. This variation likely reflects intraspecies genomic diversity, differences in sequencing depth, or annotation pipelines (22).

**Figure 1.**
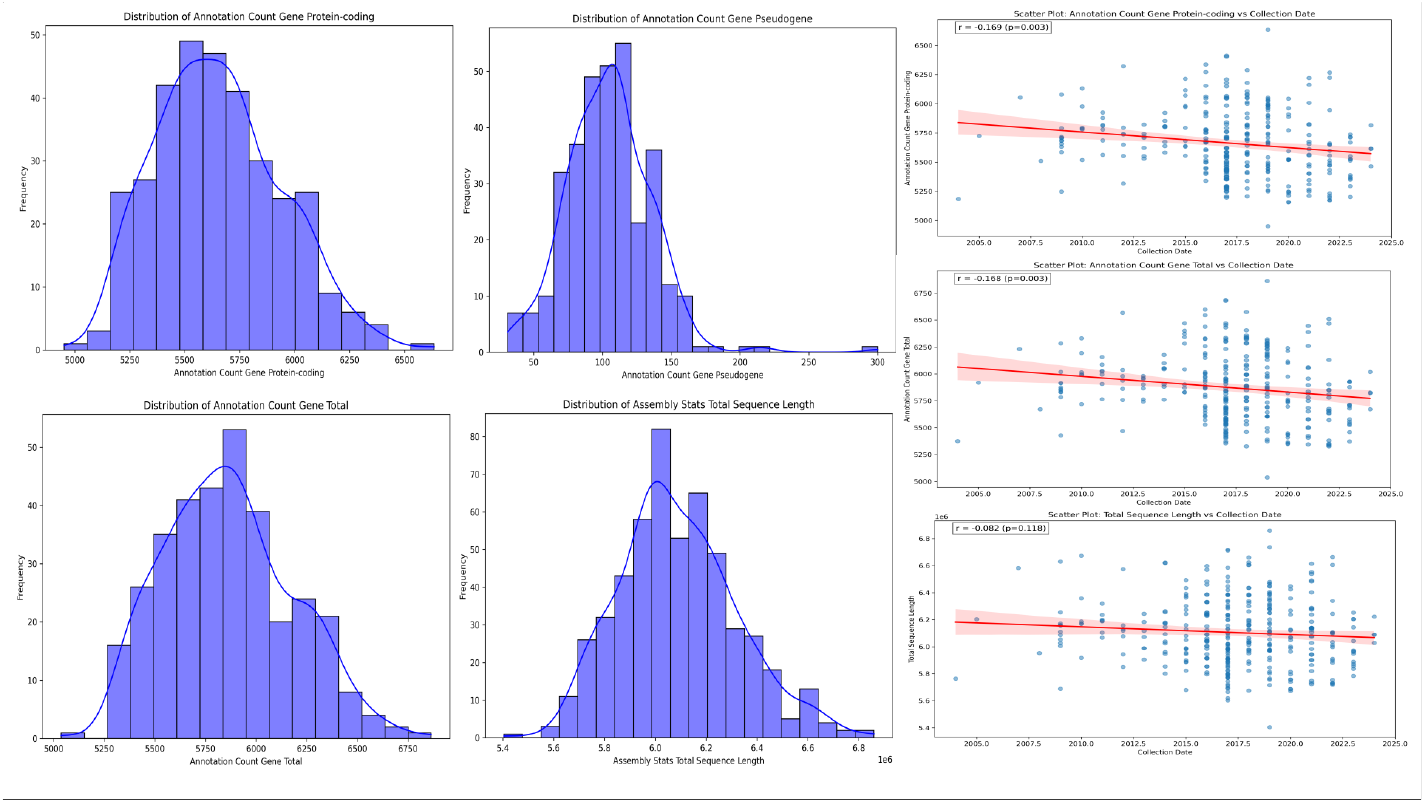
Distribution of different parameters of *K. oxycota* genomes.

FetchM’s metadata enrichment feature revealed critical insights into sample context. Geographic location was missing for 23.9% of the genomes. The most represented countries were Poland (17.6%), Switzerland (14.0%), USA (13.6%), China (7.1%), and Australia (5.5%) (Figure 2). Metadata on host showed that 64.4% of isolates originated from *Homo sapiens*, underscoring the species’ clinical importance, while 27.3% of records lacked host data. Other isolates came from diverse sources, including domestic animals, environmental reservoirs, and less common origins such as kidney bean, cricket, and the fish *Mystus tengara*.

**Figure 2.**
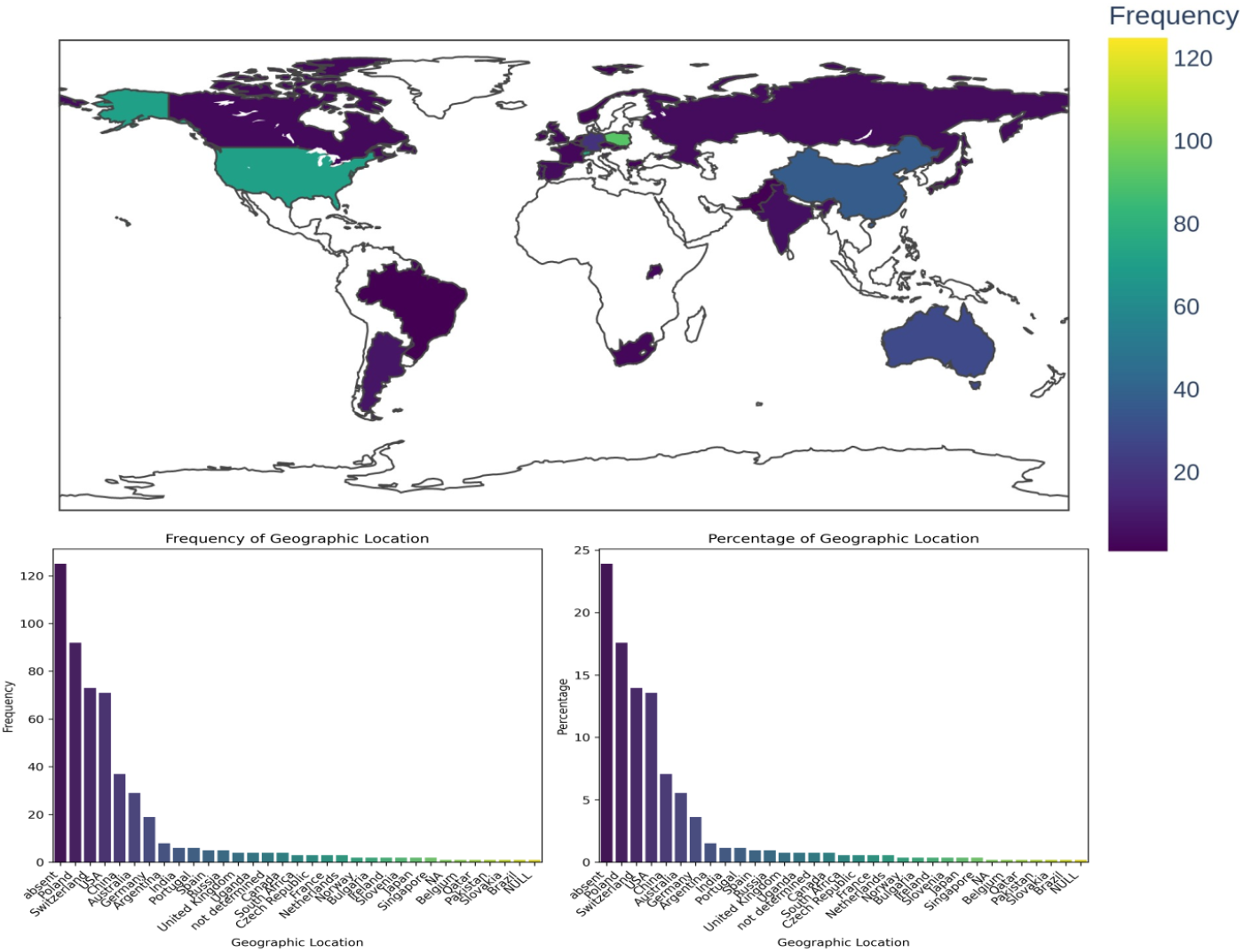
Distribution of geographic sources of *K. oxycota* genomes

Regarding collection dates, 29.6% of records were missing the information, but most isolates were collected between 2015 and 2023, with a peak in 2017 (Figure 3). Some genomes dated as far back as 2004, while others were collected as recently as 2024, highlighting FetchM’s ability to retrieve and consolidate the most current and historical data (23).

**Figure 3.**
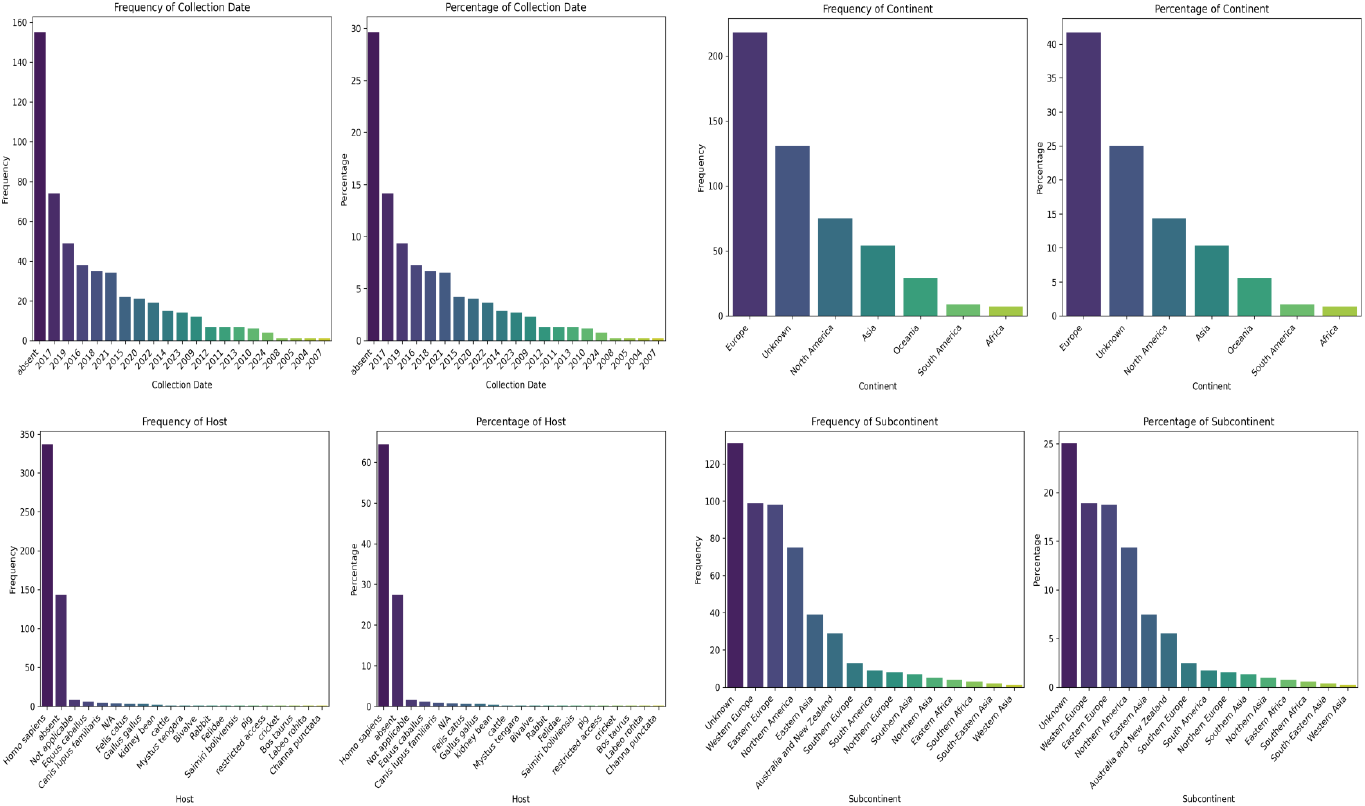
Distribution of *K. oxycota* genomes based on host, collection year, continent, and subcontinent.

Beyond merging fragmented metadata, FetchM enables custom filtering of genomes by parameters like host, country, and year of isolation. This is especially beneficial for comparative genomics, phylogeography, and population-level epidemiology, where metadata-driven stratification is key (24). Additionally, FetchM provides summary statistics and preliminary visualizations for quality inspection and exploratory data analysis using built-in plotting tools (25).

Overall, FetchM offers a streamlined solution for integrating genomic and contextual data from NCBI Genome and BioSample into a single, high-quality, customizable dataset. While dependent on the completeness of public records, its modular design, automation, and export options make it a robust asset for microbial genomics and surveillance studies (26).

## Author contributions

T.A.A. conceptualized the study, supervised the research, and developed the computational tool.

T.A.A. drafted the original manuscript, critically revised subsequent versions, and approved the final submitted draft.

## Data and code availability

FetchM is available as an open-source tool at:

https://github.com/Tasnimul-Arabi-Anik/FetchM. It can also be downloaded as a PyPI package.

## References

1. Sayers EW, Cavanaugh M, Clark K, et al. GenBank. Nucleic Acids Res. 2022;50(D1):D161–D164.

2. Barrett T, Clark K, Gevorgyan R, et al. BioProject and BioSample databases at NCBI: facilitating capture and organization of metadata. Nucleic Acids Res. 2012;40(Database issue):D57–D63.

3. Benson DA, Cavanaugh M, Clark K, et al. GenBank. Nucleic Acids Res. 2018;46(D1):D41–D47.

4. Tettelin H, Riley D, Cattuto C, Medini D. Comparative genomics: the bacterial pan-genome. Curr Opin Microbiol. 2008;11(5):472–477.

5. Federhen S. The NCBI Taxonomy database. Nucleic Acids Res. 2012;40(Database issue):D136–D143.

6. Kans J. Entrez Programming Utilities Help [Internet]. Bethesda (MD): National Center for Biotechnology Information (US); 2023.

7. Sayers EW, Bolton EE, Brister JR, et al. Database resources of the National Center for Biotechnology Information. Nucleic Acids Res. 2021;49(D1):D10–D17.

8. Cock PJA, Antao T, Chang JT, et al. Biopython: freely available Python tools for computational molecular biology and bioinformatics. Bioinformatics. 2009;25(11):1422–1423.

9. Shah N, Nair A, Sinha A, et al. pysradb: a Python package to query next-generation sequencing metadata and data from NCBI Sequence Read Archive. F1000Res. 2021;10:294.

10. Wickham H. ggplot2: Elegant Graphics for Data Analysis. Springer-Verlag New York; 2016.

11. Parks DH, Imelfort M, Skennerton CT, Hugenholtz P, Tyson GW. CheckM: assessing the quality of microbial genomes recovered from isolates, single cells, and metagenomes. Genome Res. 2015;25(7):1043–1055.

12. Cock PJ, Antao T, Chang JT, et al. Biopython: freely available Python tools for computational molecular biology and bioinformatics. Bioinformatics. 2009;25(11):1422–3.

13. Parks DH, Imelfort M, Skennerton CT, Hugenholtz P, Tyson GW. CheckM: assessing the quality of microbial genomes recovered from isolates, single cells, and metagenomes. Genome Res. 2015;25(7):1043–1043.

14. McKinney W. Data structures for statistical computing in Python. In: van der Walt S, Millman J, editors. Proceedings of the 9th Python in Science Conference. 2010. p. 51–6.

15. Hunter JD. Matplotlib: A 2D graphics environment. Comput Sci Eng. 2007;9(3):90–90.

16. Waskom ML. Seaborn: statistical data visualization. J Open Source Softw. 2021;6(60):3021.

17. Plotly Technologies Inc. Collaborative data science. Montréal, QC: Plotly Technologies Inc.; 2015.

18. Page AJ, Cummins CA, Hunt M, et al. Roary: rapid large-scale prokaryote pan genome analysis. Bioinformatics. 2015;31(22):3691–3.

19. Tonkin-Hill G, MacAlasdair N, Ruis C, et al. Producing polished prokaryotic pangenomes with the Panaroo pipeline. Genome Biol. 2020;21(1):180.

20. Eren AM, Esen ÖC, Quince C, et al. Anvi’o: an advanced analysis and visualization platform for ‘omics data. PeerJ. 2015;3:e1319.

21. NCBI Entrez Programming Utilities Help [Internet]. Bethesda (MD): National Center for Biotechnology Information (US); [cited 2025 Apr 7]. Available from: https://www.ncbi.nlm.nih.gov/books/NBK25501/

22. Touchon M, Hoede C, Tenaillon O, et al. Organised genome dynamics in the Escherichia coli species results in highly diverse adaptive paths. PLoS Genet. 2009;5(1):e1000344.

23. Li W, Jaroszewski L, Godzik A. Clustering of highly homologous sequences to reduce the size of large protein databases. Bioinformatics. 2001;17(3):282–3.

24. Didelot X, Bowden R, Wilson DJ, et al. Transforming clinical microbiology with bacterial genome sequencing. Nat Rev Genet. 2012;13(9):601–12.

25. Hunter JD. Matplotlib: A 2D graphics environment. Comput Sci Eng. 2007;9(3):90–5.

26. Cury J, Touchon M, Rocha EPC. Integrative and conjugative elements and their hosts: composition, distribution and organization. Nucleic Acids Res. 2017;45(15):8943–56.

